# Generation of a Synthetic Single Domain Antibody Library for Radiopharmaceutical Ligand Discovery

**DOI:** 10.1101/2025.05.14.654066

**Authors:** Lucinda Hall, Rachael Guenter, Rodrigo Queiroz, Amber Jackson, Yuvasri Golivi, Jasmine Watts, Yujun Zhang, Luke Rathbun, J. Bart Rose, Benjamin M. Larimer

**Affiliations:** University of Alabama at Birmingham, Department of Radiology; University of Alabama at Birmingham, Department of Surgery; O’Neal Comprehensive Cancer Center, University of Alabama Birmingham

**Author notes:** Corresponding author: Benjamin M. Larimer.

## Abstract

Single domain antibodies, often known as nanobodies, are versatile molecules with therapeutic and diagnostic applications, but they are primarily developed through immunization of camelids. This approach is not scalable by automation, not effective for non-immunogenic or toxic antigens, and prevents the use of modified scaffolds for altered pharmacokinetic properties. Synthetic libraries allow for pre-selection of a single domain framework tailored to its intended downstream use. One area of interest for these biologic vectors is radiopharmaceuticals. Ideal radiopharmaceutical pharmacokinetic properties differ from most traditional therapeutics, as short plasma circulation and rapid kidney clearance are necessary to avoid dose-limiting organ radiation. Although there are a growing number of nanobody radiopharmaceuticals in clinical trials, their frameworks and corresponding pharmacokinetic properties vary. One potential method for improving the development of novel single domain antibody radiopharmaceuticals is through synthetic libraries based on nanobodies with proven clinically acceptable pharmacokinetics. We developed a modular synthetic nanobody phage display vector based on the scaffold of the 2Rs15d nanobody that allows for manipulation of the binding and framework regions. Using this vector, we created a library of nanobodies with a randomized CDR2 containing over 1.7×10^6^ unique sequences/µL. As a proof-of-concept, we panned the library for nanobodies binding calreticulin (CALR), a protein critical in immunogenic cell death. One isolated clone, Cal3, has a measured affinity of 140 nM for CALR and is cross-reactive with mouse and human CALR. Using positron emission tomography (PET) imaging, the radiolabeled ^64^Cu-NOTA-Cal3 demonstrated CALR binding *in vivo*, representing the first reported synthetic nanobody characterized by PET imaging. This study demonstrates the feasibility of building and panning synthetic libraries for high-affinity radiopharmaceutical nanobodies as an alternative to immunized camelid libraries.

## INTRODUCTION

Radiopharmaceuticals, including positron emission tomography (PET) imaging agents, have emerged as a valuable clinical strategy for characterizing biological processes and quantifying tumor antigens. However, the development of PET tracers for biomarkers of interest has been historically limited. Single domain antibodies, often referred to as nanobodies, represent a promising class of biologic PET ligands. Derived from the variable domain of heavy-chain-only antibodies found in camelids, they are compact and versatile antibody fragments that have emerged as biologic tools with significant implications for both imaging and therapeutic purposes [1]. Their unique structural features, characterized by a single monomeric domain comprised of a rigid scaffold and defined binding regions, endow nanobodies with high stability, solubility, and binding specificity to a target of interest. In radiopharmaceutical applications, nanobodies exhibit advantages over traditional immunoglobulins including improved tissue penetration and rapid clearance, making them attractive candidates for non-invasive diagnostic agents [2].

Nanobodies are typically discovered through immunization to a target antigen of interest [3]. While immunization has proven to be a valuable method for nanobody discovery, it has limitations. Non-immunogenic antigens, molecules with structures that are not stable or unable to be formulated for immunization, and inability to direct selectivity towards specific epitopes all represent challenges to existing immunization strategies [4]. Synthetic phage display libraries can directly address these limitations. Additionally, the controlled and systematic design of synthetic libraries allows for the incorporation of specific structural features, such as varied complementarity-determining region (CDR) lengths and amino acid compositions, enhancing the range of desired nanobody properties [5]. Rational engineering enables the selection of nanobodies against targets that may be challenging or impractical to use in immunization strategies, including toxic or non-immunogenic antigens. Moreover, synthetic libraries are not limited by species-specific immune responses, making them versatile tools for targeting a diverse array of antigens [6].

Here we report the development of a novel synthetic nanobody phage display library. The framework of this library is based on the 2Rs15d nanobody, a well characterized ligand that is currently in clinical trials for PET imaging [7, 8]. The 2Rs15d nanobody, initially identified in 2011 by Van Eckyen et al. through biopanning a HER2 immunized camelid phage display library, has been utilized in a number of studies [9]. It has been researched both in preclinical and clinical contexts, serving as a PET imaging agent and targeted radiotherapy for HER2^+^ cancers [10-13]. Furthermore, a structural analysis of its interaction with HER2 has been conducted [12, 14, 15]. Taken together, these data support the use of 2Rs15d as a scaffold for synthetic library design.

To demonstrate the utility of a 2Rs15d-based synthetic nanobody library for PET imaging agent generation, selections for nanobodies specific to calreticulin (CALR) were conducted. Calreticulin, a multifunctional endoplasmic reticulum-resident protein, has gained increasing significance as a biomarker in cancer therapy [16]. Its role extends beyond its classical function in calcium homeostasis and protein folding, as emerging research suggests that altered calreticulin expression is associated with various cancers [17]. Importantly, the involvement of calreticulin in immune response modulation and its expression during immunogenic cell death, makes it a strong candidate biomarker for interrogating the tumor microenvironment [18]. A non-invasive biomarker for calreticulin could be used to assess the effectiveness of specific therapies in inducing immunogenic cell death, act as guide for combination immunotherapies, or even serve as a cancer-specific therapeutic target. To date, only one peptide-based PET imaging agent has been described although non-imaging agents including a monobody and nanobody have also been reported [19-21]. While this peptide has been demonstrated to detect increases in calreticulin following doxorubicin treatment, its micromolar affinity may limit clinical utility.

In this study, we have panned our synthetic nanobody library for a nanobody with affinity and specificity for CALR, demonstrate its cross reactivity with mouse and human CALR, and assess its *in vivo* detection of CALR via PET imaging. These data demonstrate a facile and rapid method for generating candidate ligands for clinical PET imaging agents.

## METHODS

### Library Cloning

A 3+3 M13 phage display library was generated by digesting a previously-reported plasmid containing the DNA sequence for a phage protein pIII protein using the KasI and NcoI (New England Biolabs) restriction enzyme cut sites [22]. Subsequently, a synthetic DNA fragment (Integrated DNA Technologies) containing a bacterial codon optimized coding sequence for the 2Rs15d nanobody was ligated into the digested vector.

A PfoI restriction site 16 bases upstream of the PelB leader sequence was altered by site directed mutagenesis using a silent mutation approach to abolish recognition by the restriction enzyme. Following PCR, the reaction was treated with KLD enzyme mix (New England Biolabs) and transformed into NEB 5-alpha competent *E. coli* (New England Biolabs). Clones were isolated and screened for successful mutagenesis by Sanger sequencing using a primer upstream of the nanobody sequence (5’ TATGACCATGATTACGCCAAGC 3’).

To randomize CDR2 residues, a single stranded oligonucleotide containing 12 NNK codon randomized residues in place of CDR2 was synthesized and amplified as a double stranded fragment via Klenow synthesis (Integrated DNA Technologies). Separately, the randomized nanobody-phagemid vector was amplified in SCS110 *E. coli* (Agilent) incapable of methylating DNA and isolated by miniprep (New England Biolabs). Three µg each of the unmethylated vector and randomized double-stranded insert was digested overnight at 37°C with 3 µL FastDigest PfoI (ThermoFisher). Following the initial digestion, the products were run on a 1% agarose gel and extracted with the Monarch® DNA Gel Extraction kit (New England Biolabs). The products were incubated overnight at 37°C with 5 units BsgI (New England Biolabs). The digested insert was desalted with a PCR clean up kit, and the digested vector was run on a 1% agarose gel and extracted as previously described. 150 ng of digested vector was ligated with a 3:1, 5:1, and 7:1 molar excess of digested insert with 1,000 units T4 DNA ligase (New England Biolabs) for 10 minutes at RT. The resulting ligation reaction was dialyzed using 0.025 µm pore size membrane filters for 4 hours. The desalted ligation reactions were transformed into Tg1 *E. coli*. Serial dilutions of each reaction were plated to calculate transformation efficiency. The ligated library was called **<u>Na</u>**nobody **<u>Li</u>**brary with randomized **<u>C</u>**DR**<u>2</u>** (NaLiC2).

### Phage Selection

To generate a capture antigen, 50 µg lyophilized recombinant human calreticulin (CALR) (Sino Biological) was reconstituted to 0.2 mg/mL in PBS and incubated with 10 molar excess of 10mM EZ-Link NHC-LC-Biotin (ThermoFisher) at 37C with shaking at 250 rpm for 1 hour. Unreacted biotin was removed by buffer exchange in an Amicon 0.5mL 10K MWCO spin column. Biotinylated CALR (Biot-CALR) was stored in TBS to neutralize any remaining unreacted biotin. Biot-CALR concentration was determined by BCA and diluted to 0.2 mg/mL. The Pierce Biotin Quantification Kit (ThermoFisher) was used to assess biotinylation of CALR.

For round 1 of selection, 5×10^12^ NaLiC2 phage was blocked in 5% BSA in Tris buffered saline plus 0.1% Tween-20 (TBST). Selections wells were prepared by adsorbing 500 ng neutravidin per well in 0.1M sodium bicarbonate (pH 9.5) to Nunc Maxisorp plates for 1 hr at room temperature (RT), followed by blocking with 5% BSA in 0.1M sodium bicarbonate (pH 9.5) overnight at 4°C. The pre-blocked library was then subtracted for neutravidin binding and nonspecific phage by incubating it in the neutravidin coated wells for 1 hr at RT with shaking at 250 rotations per minute (RPM). The subtracted library was then transferred to a microcentrifuge tube and incubated with 1 µg biotinylated CALR (Sino Biological) with shaking at 250 RPM. The phage were transferred to a neutravidin well to capture phage bound to target by shaking at 250 RPM for 1 hour at RT. The captured phage were eluted with 100 µL 0.05% trypsin-EDTA (Gibco) shaking at 37°C, 250 rpm for 5 minutes and neutralized with 100 µL Pierce EDTA-free protease inhibitor tablet (ThermoFisher) dissolved in 10 mL TBS. Eluted phage were titered and amplified according to previously described protocols [22]. For round 2, selection proceeded as in round 1 with the following exception: target binding occurred with 1 µg unbiotinylated CALR adsorbed directly to the polystyrene plate. Round 3 of selection proceeded identically to round 1. Phagemid were isolated from each round of selection for deep sequencing to monitor for individual clone enrichment.

### Sequencing and Analysis

The CDR2 of the starting library and eluted phage from each round of selection was amplified via PCR of primers flanking CDR2 (5’ TATCGGCAGAGTCCTGGAAGAGAG 3’ and 5’ GTTTCCAAGTTGTAACACACTGCAC 3’ and utilized for next generation sequencing (NGS) at the Heflin Genomics Core at the University of Alabama at Birmingham (Birmingham, AL, USA). A Python program connected to a MySQL database was used for analysis of NGS output. Analysis was performed by splicing the parent sequence at the first ‘N’ and last ‘K’ to create ‘heads’ and ‘tails’ sequences. The uploaded sequences were initially searched for the ‘heads’ sequence, and those that match were sorted into a separate list for searching for the ‘tails’ sequence. The sequences that had both were determined to be useful and were trimmed to retain only the CDR sequence. Following translation using a hardcoded codon table, reads per million (RPM) was calculated as a normalization parameter. This was accomplished by dividing the raw count of sequences by the total number of sequences and scaled to a million. Next, a table was created using the input experiment name with 3 columns: sequence, number of repeats, and RPM. The analysis further involved iterating through sequences to count the occurrences of each amino acid at every position. These values were stored in dictionaries, with the position as the key and frequency as the value. Frequencies were computed by dividing the number of appearances by the total number of sequences, multiplied by the number of codons resulting in that amino acid, and then multiplied by 100 to obtain a percentage. These dictionaries, containing positions and frequencies for each amino acid, were organized into 12×20 matrices (representing 12 positions for 20 amino acids) for utilization in NumPy arrays, facilitating visualization through heatmaps.

### Nanobody Cloning, Expression, and Purification

The codon-optimized CDR2 DNA sequences corresponding to the most enriched amino acid sequences (Cal1: STHVSALLYSKG, Cal2: GCLAVCPHRKT, Cal3: SVGDCMAGSSPN) for the selected clones were ligated into the 2Rs15d nanobody sequence. A modified pET28a (+) vector was used for cytoplasmic expression of the screened nanobodies. Additionally, the vector included a C-terminal Factor Xa cleavage site, FLAG, and 6XHIS tags. 50 ng of digested vector was ligated with a 3:1, 5:1, and 7:1 molar excess of digested insert with 400 units T4 DNA ligase (New England Biolabs) for 10 minutes at RT. The resulting ligation reaction was transformed into NEB 5-alpha competent *E. coli* (New England Biolabs). The plasmid was isolated from spots from each ligation reaction and sequenced from the T7 promoter upstream of the nanobody. Plasmids containing successfully ligated clones were transformed into T7 Shuffle competent *E. coli* (New England Biolabs).

Overnight cultures of Nb-pET28 T7 Shuffle were prepared in 50 mL LB supplemented with 50 µg/mL kanamycin sulfate and 0.5% glucose. Overnight culture was then diluted to OD_600_ 0.05-0.1 in 400 mL LB supplemented with the same concentration of kanamycin sulfate and 0.01% glucose and incubated until OD_600_ reached 0.6-0.8. IPTG was added at a final concentration of 0.01 mM, and the induced cultures were grown for 16-20 hours at 23°C with agitation at 250 RPM. Induced cultures were pelleted by centrifugation at 3,000 x g for 20 minutes. Pellets were frozen, thawed, and resuspended in 20 mL B-PER Complete Bacterial Protein Extration Reagent supplemented with protease inhibitors (ThermoFisher) rocked at 23°C for 1 hour. The solution was pelleted by centrifugation at 10,000 RPM for 15 minutes and the supernatant collected. Protein extract was immediately purified.

Nanobodies were purified by immobilized metal affinity chromatography (IMAC). Protein extract was equilibrated to 10mM Tris (pH 8), 200 mM NaCl, and 10mM Imidazole and added to Ni-NTA Agarose (ThermoFisher) washed in binding buffer composed of 10mM Tris (pH 8), 200 mM NaCl, 10 mM imidazole, and 0.05% TritonX. The extract-agarose mixture was rocked overnight at 4°C. Agarose was pelleted and flow through was removed. The pellet was washed 3 times in wash buffer identical to binding buffer with 20 mM imidazole. Nbs were eluted by 3 washes with elution buffer identical to binding/wash buffer with 250 mM imidazole. Fractions containing Nb were confirmed by SDS-PAGE. Cal3 used for *in vivo* PET imaging studies underwent cleavage of detection and purification tags. Cal3 was exchanged into a buffer containing 100 mM NaCl, 20 mM Tris (pH 8), and 2 mM CaCl2 using a 0.5 mL 3K spin column (Millipore). After exchange, 10 µg Factor Xa Protease (NEB) was added, and the sample was shaken at 250 RPM at 23°C for 2-4 hours.

Size Exclusion Chromatography (SEC) was conducted using the GE AKTA FPLC. Nanobody elutions resulting from IMAC purification were loaded onto the Superdex Increase 75 10/300 GL column (Cytiva Life Sciences) after the column was prepared by washing with 2 column volumes (48 mL total) of High-Performance Liquid Chromatography (HPLC) grade water and one column volume (24 mL) of Tris Buffered Saline (TBS). Eluent A was HPLC-grade water and eluent B was TBS. The FPLC method is as follows: inject sample onto Superdex 75 Increased 10/300 GL column washed with TBS. Elute with 100% eluent B (TBS) for 1.5 column volumes at 0.8 mL/min flow rate. Fractions containing nanobody were identified via the UV peaks on the FPLC and verified via running SDS-PAGE gels. The nanobody-containing fractions were then buffer-exchanged using a 0.5 mL 3K spin column (Millipore) and washed with metal-free phosphate buffer saline (PBS). Nanobody concentration was determined by BCA.

### ELISA Screening

100 ng of recombinant protein per well in 0.1 M sodium bicarbonate (pH 9.5) was immobilized on Maxisorp plates (ThermoFisher) and blocked with 1% milk in 0.1 M (pH 9.5) sodium bicarbonate. Nanobody dilutions from 2.5 mM to 100 nM were added to wells and incubated with shaking at RT for 1 hour. The plate was washed 15 times with 300 µL TBST. 1:1000 dilution of anti-FLAG-HRP antibody (Millipore Sigma) was added to each well and incubated for 1 hour at RT with shaking. The plate was washed 10 times with TBST. TMB (Millipore Sigma) was added to each well and covered with foil and shaken at RT for 15 minutes. Reaction was quenched with 2 M sulfuric acid and read at 450 nm on a ThermoScientific Varioskan LUX plate reader. Binding curves were calculated using GraphPad Prism 10.4.1. The curves were calculated using nonlinear regression analysis of one site total binding and one site total and nonspecific binding.

### Biolayer Interferometry (BLI) Analysis

Biolayer interferometry (BLI) was performed using an Octet RED (ForteBio) controlled by Octet DataAnalysis software (version 9.0.0.14). Streptavidin (SA) biosensors (Sartorius, 18-5020) were hydrated in TBST for at least 10 minutes before use.

For ligand immobilization, biotinylated CALR (as described above) was diluted to 10 µg/mL in TBST and loaded onto the biosensors for 600 s at 1000 rpm. After loading, biosensors were washed in TBST to remove unbound ligand. Binding kinetics were assessed by exposing the functionalized biosensors to a series of Cal3 nanobody (Nb) concentrations ranging from 23.8 nM to 22.7 µM in a 3.33-fold dilution series. Association and dissociation were monitored for 300 s each at room temperature (RT). A reference biosensor was used to correct for non-specific binding.

Data were analyzed using Octet DataAnalysis software (version 9.0.0.14), and kinetic parameters (k_a_, k_d_, and K_D_) were determined using a 1:1 binding model with global fit and steady-state analysis. Data were baseline-corrected using reference well subtraction, and negative controls included a buffer-only reference well.

### CALR Sequence Conservation Analysis

The amino acid sequences of human calreticulin (P27797) and mouse calreticulin (P14211) were obtained from UniProtKB and aligned with Smith-Waterman local alignment using Snapgene software (www.snapgene.com) [23].

### Flow Cytometry

PDAC cells (2838c3 cell line) were cultured in DMEM media supplemented with 10% fetal bovine serum and 1% penicillin-streptamycin and grown to 70% confluency, followed by treatment with either 100nM doxorubicin or 250µM H2O2 for 24h to induce surface CALR. DMSO was used as vehicle control. After treatment, cells were harvested and stained with UV Zombie Dye (Biolegend 423108) at a 1:250 dilution as a live-dead stain. After washing, cells were incubated with Cal3 nanobody at 0.1 mg/mL for 30 minutes at 4°C in the dark. After nanobody incubation, cells were fixed with 4% PFA, followed by incubation with a rabbit anti-FLAG primary antibody (CST #23685) at a 1:200 dilution for 30 minutes at 4°C in the dark. An anti-rabbit-A647 secondary antibody at a 1:2450 dilution was used. Samples and appropriate compensation controls were analyzed using BD FACSymphony A5 SE channels UV446, R660, and B537.

### Bioconjugation and Radiolabeling

Nanobodies were conjugated with p-SCN-Bn-NOTA (Macrocyclics) in a 10-fold molar excess of chelator. Reaction occurred for 1 hour in metal-free HEPES buffer (pH 8) at 37°C with shaking at 350 rpm. The reaction was quenched with metal-free 0.5M Tris-HCl (pH 8) and the conjugated Nbs were buffer exchanged into metal-free PBS.

^64^Cu was obtained from the UAB Cyclotron Facility (Birmingham, AL, USA). It was adjusted to pH 6 with metal-free 1M sodium acetate (pH 6) and 185 MBq was added to 1 mg conjugated Nb. The reaction was incubated at 37°C for 1 hour with shaking at 500 rpm. Free ^64^Cu was removed by filtration through spin column. Radiolabeled Nb was diluted in sterile saline to 1 mCi/mL.

Radiochemical purity was assessed using radio-thin layer chromatography (radio-TLC) on silica chromatography paper (Agilent, SIG001). 1 µL (∼40 kBq) of the sample was spotted and developed in 50 mM EDTA until the solvent front reached 150 mm. Radioactivity distribution was immediately measured using an AR-2000 TLC scanner (Eckert & Ziegler) equipped with a gas flow proportional detector. Retention factors (R_f_) were calculated as the ratio of compound migration to solvent front migration. Radiochemical purity was determined as the percentage of radioactivity at R_f_ ∼0.12 relative to total detected activity.

### Imaging and Analysis

The animal study protocol (IACUC-22619) was approved by the Institutional Animal Care and Use Committee of The University of Alabama at Birmingham in August 2022. Male C57/BL6 mice were subcutaneously implanted with Matrigel plugs containing either recombinant CALR or BSA at a final concentration of 3.7 µM injected on opposite shoulders immediately before radiotracer injection. Mice were injected intravenously with ∼80 µCi ^64^Cu-NOTA-Cal3. Under anesthesia, mice were imaged 1-hour post-injection on a Sofie GNEXT PET/CT scanner for 20 minutes followed by a 5-minute CT at 80 kVp. Acquired PET images were analyzed using VivoQuant (Invicro) software. Regions of interest (ROIs) were drawn using anatomic guidance of the CALR and BSA plugs, heart, kidneys, and liver. The lower ventricles of the heart were used as a measure of blood signal. The signal was used to calculate standardized uptake values (SUVs). SUVs of CALR and BSA plugs from PET imaging were assessed using unpaired t-tests (Graphpad Prism 10.0.2).

### Immunohistochemistry

Tumors were isolated from mice treated with doxorubicin as described in PET imaging studies and fixed in 4% paraformaldehyde, paraffin embedded, and sliced in 5 µm sections. The slides were baked overnight at 60°C prior to deparaffinization and hydration of tissue sections. Sections were washed in xylene and decreasing concentrations of ethanol ranging from 100% to 75%, followed by dH_2_O washes. For antigen retrieval, sections were boiled in citrate buffer at pH 6.0 and then cooled to room temperature. Slides were incubated in 3% H2O2 in TBS for 10 minutes followed by several washes with TBS. Tissue sections were outlined with a hydrophobic pen and blocked in 3% serum in 0.3% Triton X-100 in TBS for 1 hour at RT. Sections being stained with Cal3 were blocked with mouse serum, and sections being stained with a control anti-CALR Ab were blocked with goat serum. Following blocking, sections were washed with TBS. Slides were stained overnight at 4°C with primary antibodies diluted in 3% serum in 0.3% Tween20 in TBS. Sections were stained with 0.25 µM Cal3 or 1:10,000 dilution of a control anti-CALR Ab (Sino Biological #13539-T60). Control secondary-only sections were blocked with primary antibody solution. Sections were washed several times with TBS prior to staining with secondary antibodies. Sections stained with Cal3 were incubated with an anti-FLAG-HRP conjugated Ab (Sigma-Aldrich #A8592) diluted 1:500 in primary antibody solution. Sections stained with the anti-CALR Ab were incubated with an anti-rabbit-HRP conjugated Ab (R&D Systems #HAF008) diluted 1:20 in primary antibody solution. Following incubation for 1 hour at RT, slides were washed in TBS. Sections were incubated with DAB chromogen for 15 minutes then dipped in hematoxylin solution, DI water, and incubated in tap water. Slides were dehydrated and cleared using sequential washes in increasing concentrations of ethanol ranging from 50% to 100%, followed by incubation in xylene. Slides were covered, dried overnight, and imaged at 40X magnification.

## RESULTS

### Generation of the Synthetic Nanobody Library NaLiC2

To select for CALR binding, CDR2 was randomized due to its lack of conserved cysteine residues involved in disulfide bonds (Figure 1A). These cysteine residues are typically present in CDR1 and CDR3 and are critical to nanobody structure. Leaving this bond intact minimizes the effect of the randomized CDR2 on the structure and functionality of the nanobody library, increasing the likelihood of isolating a highly specific binder. The resulting nanobody library is referred to as Nanobody Library of randomized CDR2, or NaLiC2.

**Figure 1.**
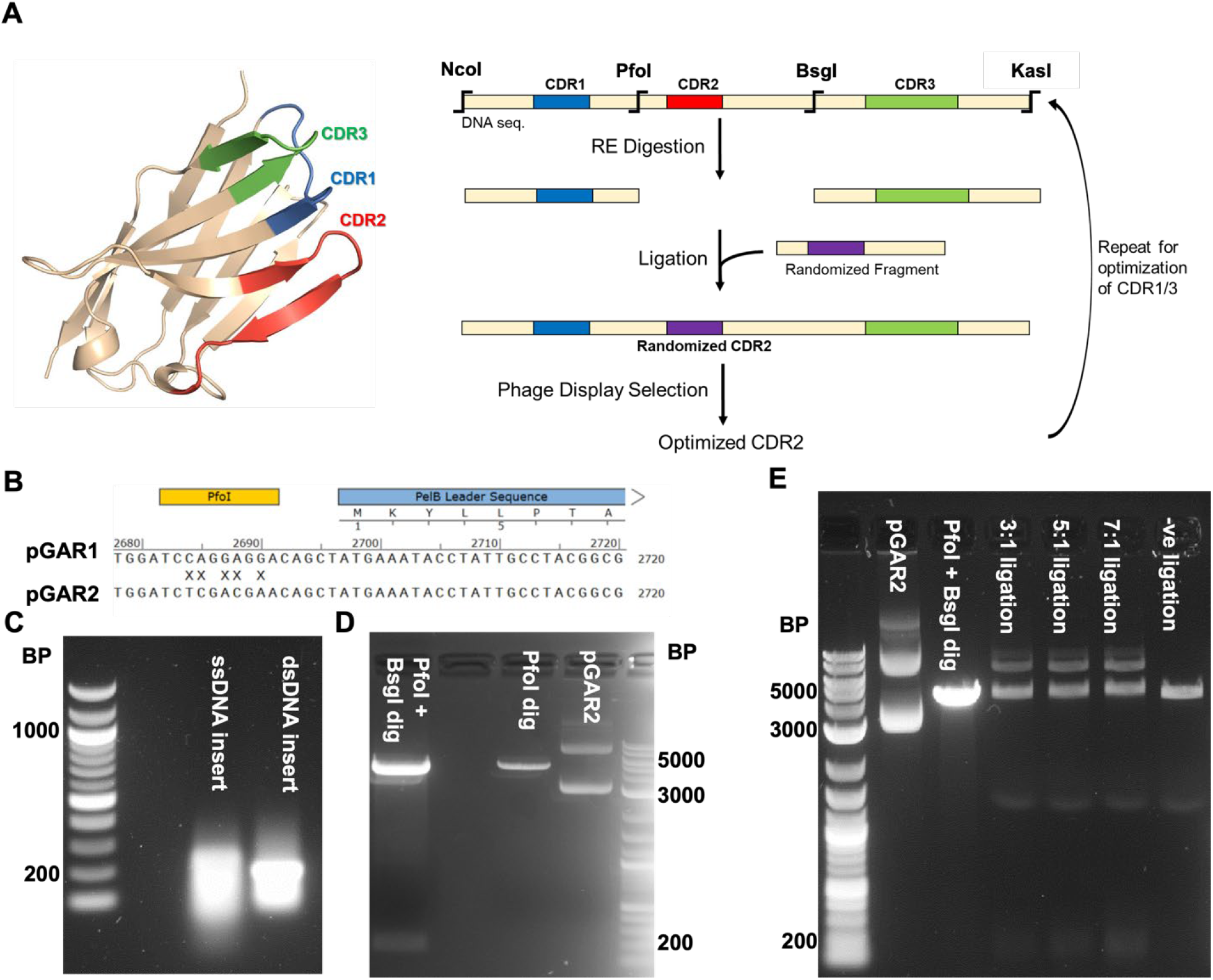
Generation of Nanobody Library with randomized CDR2 (NaLiC2). A) Schematic of the strategy used to randomize regions of the Nb phage library. B) Sequencing alignment showing mutation strategy for removal of the PfoI restriction site upstream of the nanobody by site directed mutagenesis. C) Agarose gel confirming the Klenow synthesis of the randomized CDR2 insert. D) Agarose gel confirming the complete PfoI and BsgI digestion of the phagemid pGAR2. E) Agarose gel confirming successful ligation of the digested randomized insert and vector.

To generate the library, a the PfoI restriction site was silently mutated using site directed mutagenesis using the strategy in Figure 1B. Klenow synthesis was confirmed by gel electrophoresis (Figure 1C), visible by the DNA appearing less diffuse after synthesis. Successful digestion of the phagemid by PfoI and BsgI was confirmed by gel electrophoresis (Figure 1D). The single cut of pGAR2 is observed by the plasmid DNA condensing to a single band of approximately 4600 base pairs. Finally, the digested insert and phagemid were ligated, and the ligation products were visible on an agarose gel (Figure 1E) in the positive ligation reactions but not the ligation reaction ran without insert.

### NaLiC2 Library Characterization

To evaluate the diversity and magnitude of the novel NaLiC2, a portion of the library was used for next generation sequencing analysis of the phagemid nanobody inserts. A heatmap of the composition of CDR2 amino acid sequences was generated (Figure 2). The library shows a slight bias for certain groups of amino acids across all CDR2 positions. These include the nonpolar residues isoleucine and methionine; polar asparagine, glutamine, and threonine; and positively charged histidine and lysine. As this heatmap was corrected for codon frequency expected for an NNK library, these data indicate some bias in DNA synthesis, PCR amplification, cloning or transformation. In addition to positional frequency, individual sequences were identified and enumerated. From the portion of the library analyzed, 1.1×10^6^ unique amino acid sequences per µL were identified, indicating 88.7% of NaLiC2 sequences were unique (Table 1). Only 0.0128% of all amino acid sequences coded for the parent 2Rs15d CDR2, indicating negligible residual parent phagemid was introduced into the library.

**Table 1.** Total amino acid sequence counts, unique amino acid sequence counts and parental sequence frequency for the NaLiC2 library.

| NaLiC2 | Per μL |
| --- | --- |
| Unique AA Sequences | 1,178,711 |
| Total AA Sequences | 1,328,845 |
| Percent Parent Sequence | 0.0128% |

**Figure 2.**
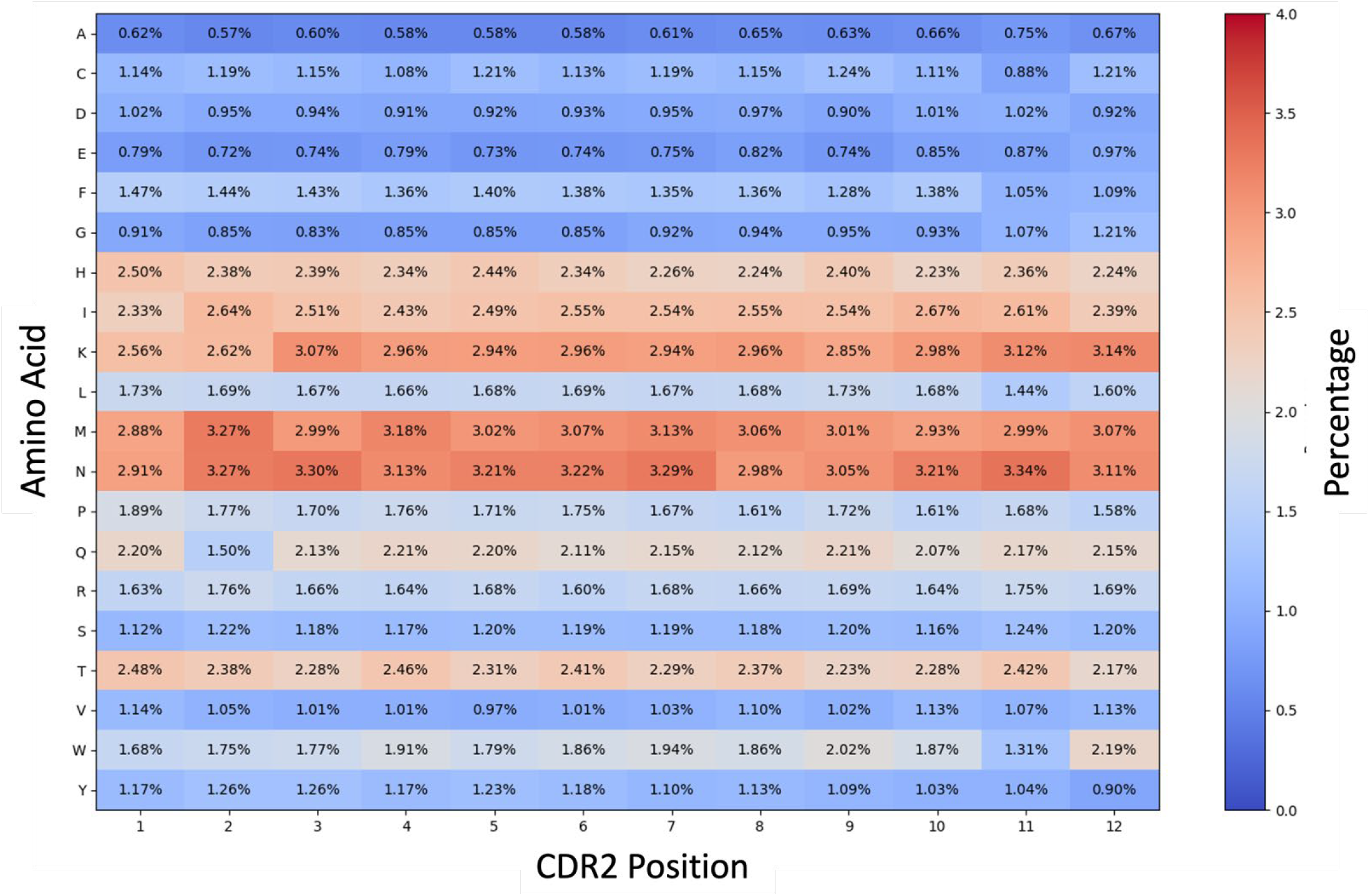
NaLiC2 positional diversity heat map. Positional frequencies for each amino acid are shown as a percentage of the total amino acids plus stop codon. Percentages shown are corrected for expected codon frequencies. Boxes are shaded using a red/blue heat map with a scale from 0-4%.

### Selection for CALR Binding Nanobodies

Following confirmation of NaLiC2 quality, three rounds of panning were performed, and each round was analyzed by quantifying phage and sequencing a portion of the library. Calculating the eluted phage as a percentage of the total used for the selection, 3.03×10^-8^% of the starting phage were recovered after round 1 of selection (Figure 3A). This recovery fell 26,000-fold during round 2, indicating an enhanced stringency in the direct target binding selection technique. 1.89×10^-10^% of phage were recovered during round 3 of selection, an increase of 62% from round 2, suggesting an amplification of CALR specific binders.

**Figure 3.**
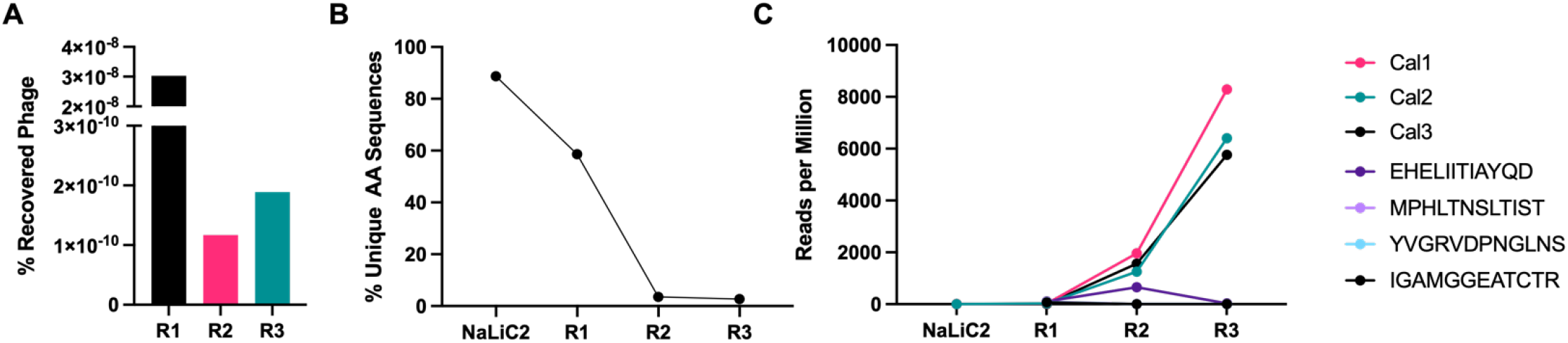
Analysis of clonal enrichment following NaLiC2.21 selection for CALR. A) Percentage of recovered phage from the total phage added for each round of selection. B) Percentage of unique amino acid sequence present in the starting NaLiC2.21 library and after each round of selection. C) The number of reads per million of each CDR2 sequence from the starting library and after each round of selection. Cal1-3 were the top 3 most enriched clones after 3 rounds of selection. The top sequences as measured as a frequency in round 1 are shown as their amino acid sequence.

Analyzing the sequence diversity after each round of selection, the most significant decrease in unique sequences occurs between round 1 and round 2 (Figure 3B). This further supports the recovered phage data that suggests that the direct target binding selection technique in round 2 is more stringent than the in-solution binding used in rounds 1 and 3. The relative stability in diversity but increase in recovered phage from rounds 2 to 3 were consistent with enrichment for target specific clones.

At an individual sequnce level, the 3 most highly enriched clones in the round 3 sequencing data were coined Cal1, Cal2, and Cal3. The abundance of these three nanobodies was quantified from the parental library through each round of selection (Figure 3C). For comparison, the frequency of the top 4 clones (EHE, MPH, YVG and IGA shown in key) from round 1 of selection were also plotted. While Cal1-3 frequencies increased each round of selection, their enrichment was the greatest between rounds 2 and 3. In contrast, the controls from round 1 increased slightly between rounds 1 and 2 of selection, but they were not present after 3 rounds of selection. The enrichment analysis of Cal1-3 was sufficient to warrant further characterization of the nanobodies outside of the context of phage display expression.

### In Vitro Characterization of Candidate CALR Nanobodies

CDR2 sequences for Cal1, Cal2, and Cal3 were cloned into a pET28a vector and expressed, and the resulting expression products were purified using the hexahistidine tag present at the C-terminus of the nanobody. Expression of Cal1 and Cal2 outside of the context of a phage particle resulted in no detectable soluble nanobody, thus these nanobodies were not investigated further. However, the Cal3 nanobody had suitable expression and could be generated in purity sufficient for subsequent analysis.

To measure the apparent affinity and specificity for CAL3 to CALR, an ELISA was utilized to compare binding to the target and control proteins including non-fat dry milk and the parental nanobody target HER2. Results from the ELISA quantified an apparent affinity of Cal3 for recombinant hCALR of 155 nM (95% CI 116.1-209.2 nM) (Figure 4A). The measured affinity for HER2 and the milk block were not quantifiable using a one-site non-linear fit. To confirm the apparent affinity of Cal3 for CALR, biolayer interferometry (BLI) was performed. Cal3 was determined to bind CALR with an affinity of 140 nM ± 44 nM.

**Figure 4.**
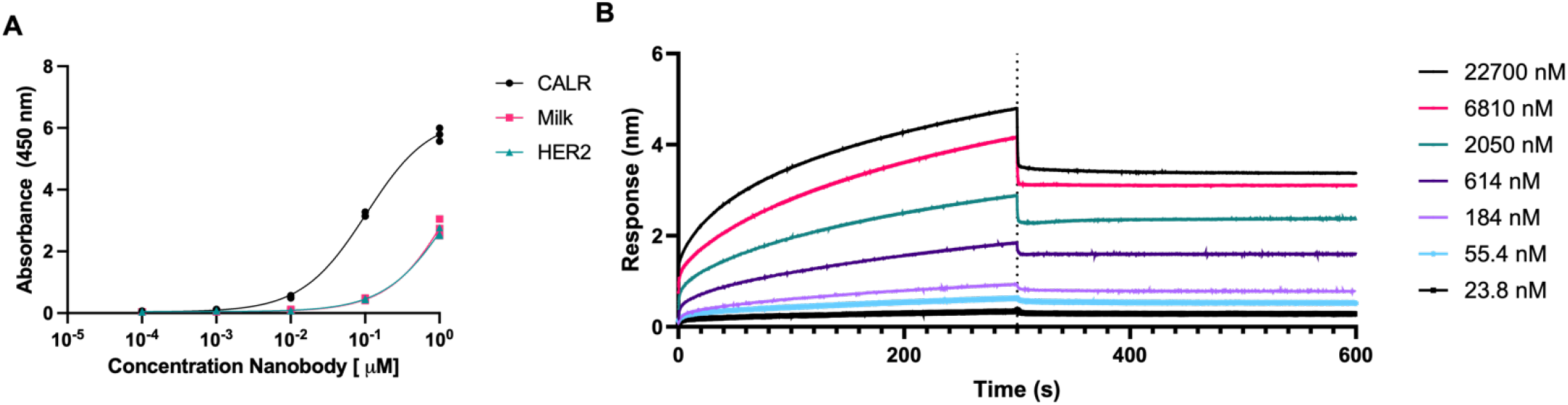
Cal3 affinity and specificity. A) Representative ELISA for Cal3 binding to CALR (Black circles), HER2 (pink squares), and 1% non-fat dry milk (teal triangles). Each point represents the mean of four individual measurements. Error bars denote standard deviation. B) Biolayer interferometry analysis of Cal3 binding to human CALR (hCALR). Individual curves representing different ligand binding concentrations from 23.8-22700 nM are depicted.

### Measurement of Binding to Secreted Cell Surface CALR

Following confirmation of high affinity and specific binding to CALR, the ability to bind extracellular CALR was next assessed. CALR translocation to the cell surface has been demonstrated using oxidative stressors including hydrogen peroxide (H_2_O_2_) and low doses of chemotherapeutics including doxorubicin. Flow cytometry was used to assess CAL3 binding to induced surface CALR in a murine pancreatic ductal adenocarcinoma (PDAC) cell line 2838c3. Although selections were performed against human CALR, an alignment of the protein sequences for both species shows that CALR is highly conserved between humans and mice with an identity of 393/417 amino acids (94.24%) and a similarity of 407/417 amino acids (97.60%). CALR translocation was stimulated by treatment with either doxorubicin or hydrogen peroxide (H_2_O_2_). Under both treatment conditions, Cal3 detected surface CALR at similar frequencies as a validated anti-CALR control antibody (Figure 4A). For further confirmation of the ability of Cal3 to bind murine cellular CALR, immunohistochemistry was conducted on sectioned 2828c3 tumors. Staining with Cal3 exhibited similar patterns observed in tissues stained with the control anti-CALR antibody. These data together indicate that Cal3 is capable of binding human and mouse CALR in both recombinant and cellular formats.

### Radiolabeling and PET Imaging of CAL3

Following confirmation of CALR binding to induced cells, the nanobody was prepared as a PET imaging agent for *in vivo* diagnostic assays. The nanobody was conjugated to the macrocyclic chelator NOTA, and radiolabeling with the positron emitting radiometal [^64^Copper]([^64^Cu]) yielded a high purity and molar activity radioligand (Supplemental Table 1). While the biological phenomenon of CALR translocation is well-established, the kinetics of its stimulation *in vivo* are not well characterized. As such, C57/BL6 mice were subcutaneously implanted with Matrigel plugs on opposite shoulders containing either hCALR or BSA as a control protein. This recombinant protein model establishes the ability of Cal3 to bind CALR *in vivo* without the confounding factors found in a tumor model. After placing the plugs, mice were injected with the [^64^Cu]-NOTA-Cal3 radioligand and scanned 1 hour post injection (Figure 6A). Visual analysis of the PET Imaging revealed moderate uptake of the probe in only the CALR containing plug, as well as uptake in the kidneys. Kidney uptake was the expected route of clearance given the molecular weight of a nanobody. Minimal uptake was visualized in other organs, including the liver. Quantification of the PET signal revealed a significantly higher accumulation of [^64^Cu]-NOTA-Cal3 in the CALR containing plugs in comparison to the control plug (0.31±0.01 CALR versus 0.12±0.04 BSA; P=0.0013) (Figure 6B). Quantification of PET signal in the blood (lower left ventricle), kidneys and liver were consistent with similar small biologic PET ligands, indicating minimal off-target organ exposure and low background signal consistent with a clinical PET tracer.

**Figure 5.**
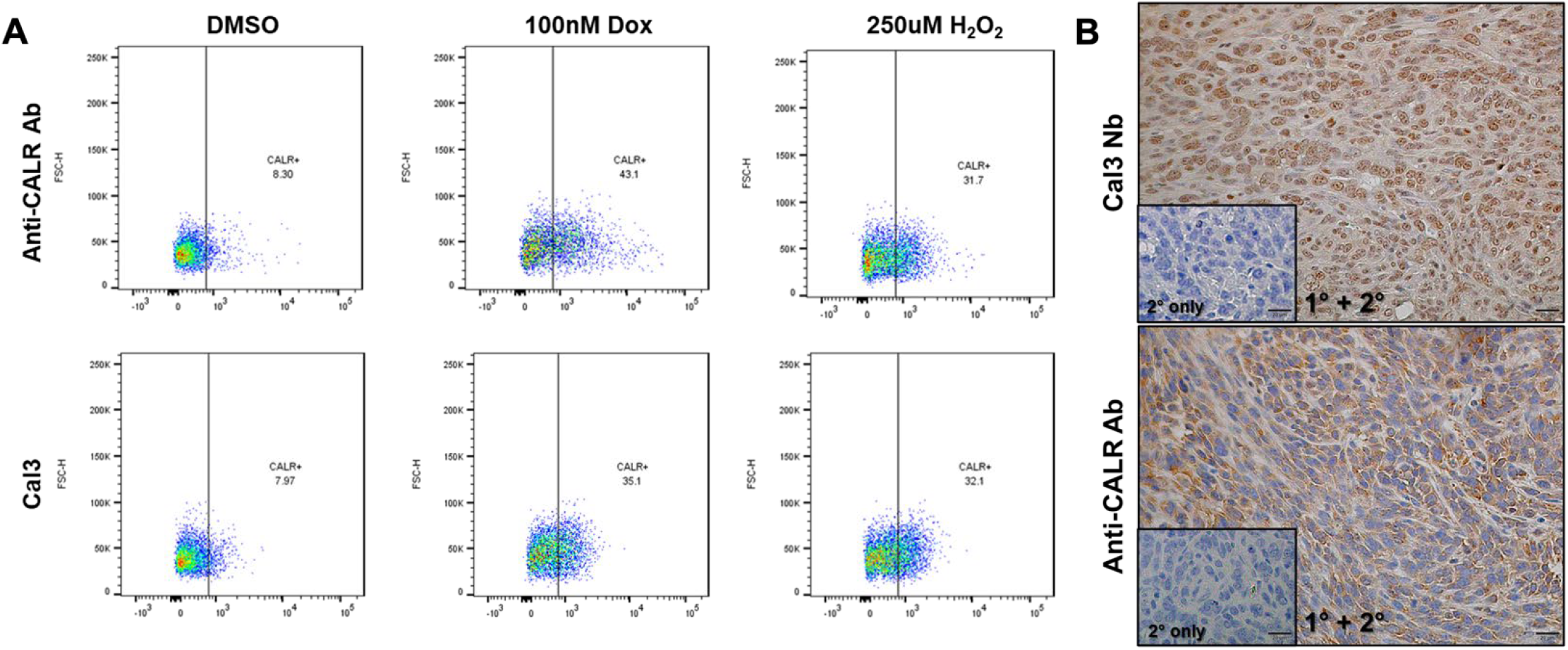
Extracellular CALR binding by CAL3. A) Representative flow cytometry scatter plots for DMSO, doxorubicin (Dox) and hydrogen peroxide (H_2_O_2_). The top row represents staining with an Anti-CALR antibody positive control, and the bottom row is stained with Cal3 and an anti-flag fluorescent secondary. The percentage of CALR+ cells are shown within each box. B) Immunohistochemistry staining of doxorubicin treated tumors with the Cal3Nb (top) and anti-CALR antibody control bottom. The inset boxes are adjacent slides stained with the appropriate secondary only controls.

**Figure 6.**
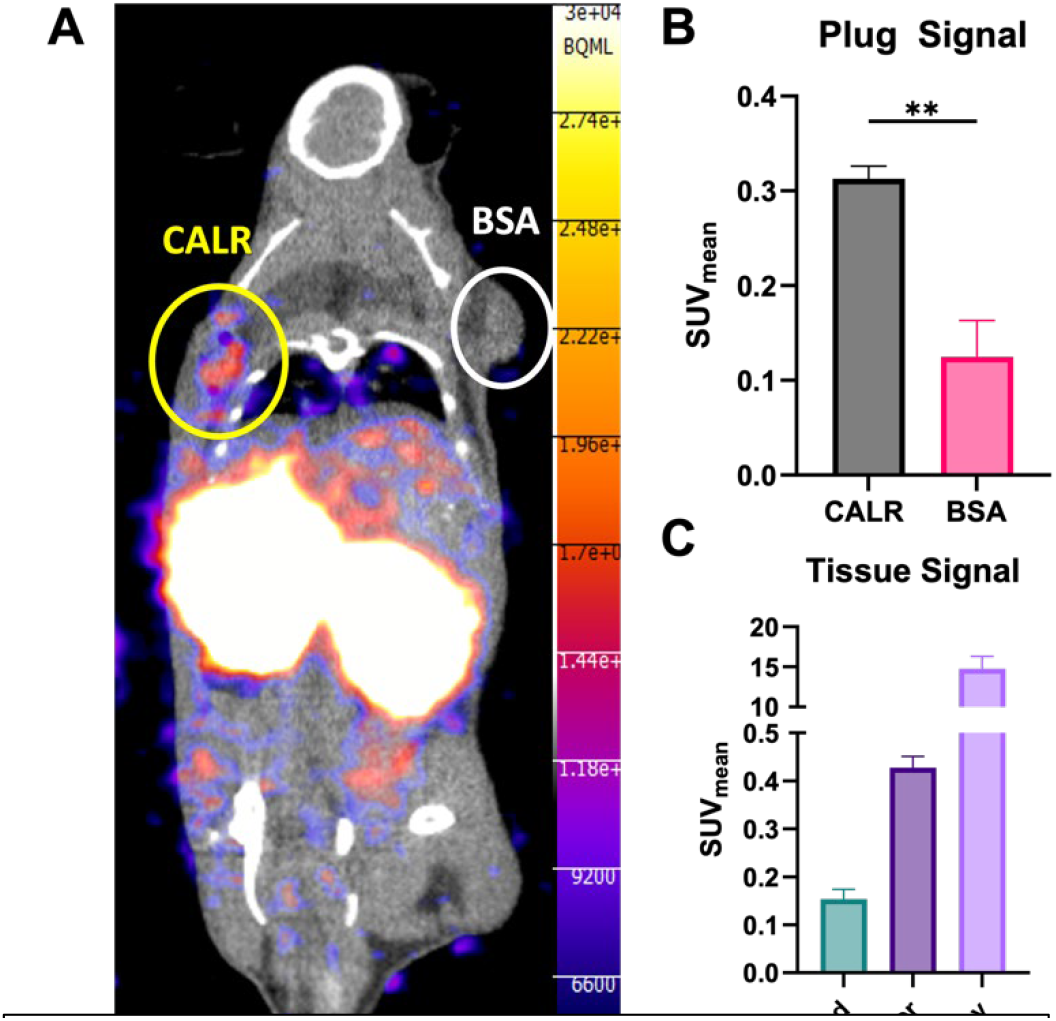
In vivo PET imaging of ^64^Cu-NOTA-Cal3 in mice bearing recombinant hCALR and BSA Matrigel plugs. A) Representative PET/CT images of ^64^Cu-NOTA-Cal3 1-hour post-injection via tail vein. Male C57/Bl6 (n=4) are bearing hCALR- or BSA-Matrigel plugs on the left and right shoulders, respectively. D) Quantification of PET signal from CALR and BSA plugs. C) Quantification of PET signal from clearance-related organs.

## DISCUSSION

Nanobodies represent a promising class of novel targeted biologics, specifically for PET diagnosis and radiopharmaceutical therapy [24-26]. A majority of nanobodies generated to date have been generated using immunization, however there are increasing reports of synthetic nanobodies [5]. In this paper, we report the first synthetic nanobody for use as a PET imaging agent. To do so, we developed a synthetic nanobody phage display library based on 2Rs15d, a clinically translated PET imaging agent [27]. The randomized CDR2 library, NaLiC2, contained over 1.7 million unique sequences per microliter, which is consistent with the reports from previous libraries [5]. From this library, we isolated Cal3, a nanobody with a CDR2 specific to calreticulin with an affinity of 140 nM. This is the second reported calreticulin nanobody and first synthetic nanobody developed for CALR [21]. Additionally, it was determined to be cross-reactive with both human and mouse calreticulin and capable of binding the cellular epitope presented during calreticulin translocation. Given its potential value as a cancer biomarker, PET imaging of calreticulin has been reported by Kim et al [19]. The ^18^F-labeled peptide KLGFFKR demonstrated significant increases in uptake post-treatment with immunogenic doses of doxorubicin. However, the affinity of the linear peptide was 1.86 µM. In comparison, we report a synthetic nanobody with 140 nM affinity and high specificity for calreticulin, with targeting of calreticulin *in vivo*.

Calreticulin represents not only a diagnostic target, but also an intriguing therapeutic target. As calreticulin is trafficked to the cell surface of cancer cells following stressors including radiation and DNA cross-linking, its expression may paradoxically increase following effective treatment [17]. This “feed-forward” target expression contrasts traditional antigens, which decrease over time. In this way, a diagnostic and therapeutic agent, such as Cal3, could be used not only labeled with a diagnostic isotope such as ^64^Cu, but subsequently labeled with ^67^Cu to be used as a therapeutic for tumors with sufficient target antigen. The non-invasive nature of PET imaging would permit interrogation of this hypothesis in future studies

One advantage of a synthetic nanobody library is that for a lead nanobody candidate such as Cal3, two additional known binding sites, CDR1 and 3, are available for subsequent function modification. For additional target avidity, a second binding site to Cal3 can be introduced using standard affinity maturation techniques. For pharmacokinetic modification, binding to albumin or a similar blood protein can be introduced. Finally, bi-specificity for immune cells or other cancer cell antigens could be imparted to increase its therapeutic index.

This study is limited in the scope of the applications that it investigates. First, we only modified one of the CDR regions, which resulted in a single function and moderate affinity nanobody. This likely limited the in vivo uptake of our PET imaging agent. Second, we limited our *in vivo* analysis to a Matrigel model of calreticulin immobilization. Given the scope of this paper was synthetic nanobody development for PET imaging, we chose to use a systematic antigen model rather than introduce additional biologic variables. Future studies should investigate previously mentioned affinity maturation and tumor imaging experiments described herein.

## CONCLUSION

Immunization provides a robust method for nanobody generation, while synthetic nanobody libraries offer a valuable and complementary approach due to their versatility, particularly for novel radiopharmaceutical development. The NaLiC2 library and Cal3 demonstrate proof-of-concept not only as viable translational agents for calreticulin but also as a pipeline for developing subsequent diagnostic and therapeutic agents.

## ACKNOWLEDGEMENTS

We thank the Cyclotron Facility at the University of Alabama at Birmingham for radionuclide production and the UAB Small Animal Imaging Facility for assistance with PET imaging and data acquisition. This work was supported by the NIH Director’s New Innovator Award (DP2CA261453-01), the NIH NIGMS Translational and Molecular Sciences Training Grant (T32GM135028), and the American Cancer Society (RSG-24-1319638-01-ET).

