## Supplemental Table 1 for "Generation of a Synthetic Single Domain Antibody Library for Radiopharmaceutical Ligand Discovery"

### SUPPLEMENTAL INFORMATION

| Radiochemical Yield | |
|---|---|
| Molar Activity | 3.43 ± 1.86 GBq/umol |
| Radiochemical Purity | 96.7 ± 3.1% |

**Supplemental Table 1** – Molar activity and radiochemical purity of $^{64}$Cu-NOTA-Cal3
